# Extracellular Matrix Proteome of Human Corneas

**DOI:** 10.1101/2025.08.08.669308

**Authors:** Khyathi Ratna Padala, PV Anusha, Mohammed M Idris, Shibu Chameettachal, Vivek Singh, Falguni Pati, Kiran Kumar Bokara

## Abstract

**BACKGROUND/OBJECTIVES:** The cornea, the transparent outer layer of the eye, is avascular and composed of three layers: a self-renewing epithelium, a transparent stroma rich in extracellular matrix (ECM) and an endothelium. This study aimed to profile ECM proteins including proteoglycans, from human decellularized extracellular matrix (hdECM) proteins using proteomics analysis.

**SUBJECTS/METHODS:** Three independent batches of lyophilised hdECM samples were subjected to proteomics analysis. Samples were processed by in-gel trypsin digestion and analysed on a Q-Exactive mass spectrometer. Protein identification and label free quantification were performed using Proteome Discoverer 2.2.0.388

**RESULTS:** Proteomic profiling identified an average of 18 proteins per batch, with 13 consistently present across all samples. Key proteins, TGF-β, keratocan, fibrillin and collagen XII, were abundant, highlighting their role in inflammation regulation, collagen organisation, and matrix remodelling. Proteoglycans such as docorins and lumican were also detected indicating their contribution to ECM structure and maintenance of corneal transparency. Network and Pathway analysis revealed involvement in ECM organization, integrin interactions, glycosaminoglycan metabolism, and collagen fibril assembly. These proteins are critical for preserving corneal architecture, modulating cell behaviour, and supporting processed such as signalling, migration, angiogenesis, and tissue repair particularly through proper collagen spacing essential for corneal transparency. This study, to the best of our knowledge, represents the first comprehensive profiling of extracellular matrix proteins from decellularised human corneas.

## Introduction

The Cornea is the transparent, avascular outermost layer of the eye that serves a protective barrier and the primary refractive surface, ensuring mechanical stability. Structurally, it comprises three major layers i) a non-keratinised, stratified squamous epithelium that acts as the first line of defence against fluid ingress and pathogens through continuous renewal of basal cells. ii) The underlying basement membrane and Bowman’s layer which support epithelial integrity and maintain stromal dehydration critical for transparency^1,3^ and iii) the stroma, which constitutes the majority of corneal volume providing transparency, shape maintenance and immune defence. The stroma is primarily composed of water and extracellular matrix (ECM) molecules, synthesized and maintained by keratocytes, which are neural crest cells responsible for ECM synthesis^2^.

The corneal stroma forms the primary structural framework of the cornea and consists of extracellular matrix rich in water, inorganic salts, proteoglycans, glycoproteins and a network of uniformly arranged, small diameter collagen fibres^3,1^. Keratocytes, the principal stromal cells, are distributed between the stromal lamellae and typically remain in inactive or resting state^4^ while maintaining ECM homeostasis. This highly ordered organization of collagen fibrils, with uniform diameter and spacing, is essential for corneal transparency by minimizing light scattering.

Decellularization is the process that removes all cellular and nuclear material from tissues or organs including immunogenic and light-scattering components, while preserving the extracellular matrix (ECM). The retained ECM provides essential structural, and biochemical cues that support cellular adhesion, proliferation, migration, and differentiation. Decellularized ECM (dECM) also maintains native growth factors, cytokines, and 3D architecture, all of which are critical for in situ tissue regeneration^6^. Decellularization can be achieved using chemical, physical, or a combination of methods^7^.

Human corneal tissues were obtained from the LV Prasad Eye Institute, Hyderabad, with appropriate donor consent. Three sets of samples comprising 44, 25 and 21 corneas respectively from both male and female donors collected in 2022 were included in the study. Corneas were decellularized using laboratory standardised protocols, lyophylised and stored until further use. The lyophilised samples were then subjected to proteomic analysis to profile extracellular matrix proteins

## Materials and Methods

### Decellularization of corneas

Human corneal tissue samples from both male and female donors collected in 2022 were obtained from the LVPEI Eye bank and classified into three sample sets (sets 1-3). The decellularization was performed according to a previously reported protocol^8^ with minor modifications. Briefly, the tissue samples were rinsed with deionized water and minced into approximately 2-3mm thickness fragments without mechanical scrapping of the epithelial or endothelial surfaces. The tissue fragments were incubated in 1.5ml sodium chloride (Himedia, India) for 72h, under continuous stirring followed by 48h wash using deionized water to remove residual salts and cellular debris. To ensure sterility 70% ethanol was added, and the suspension was stirred for 2 hours. The decellularized corneal tissues which turned visibly white appearance were subsequently lyophilised (n = 3).

*Sample preparation of decellularized corneal ECM for mass spectrometry analysis:* Lyophilised hdECM samples were solubilised in 6M Urea containing 10% SDS and processed for proteomics analysis. Approximately 50µg of protein from each sample was resolved on a 12% SDS-gel until the dye front migrated ∼2 cm. Each lane was divided into three fractions for ingel trypsin digestion. Gel bands were exised, destained and washed with 70% ammonium bicarbonate (ABC) and 30% acetonitrile (ACN). Proteins were reduced with 10 Mm dithioethriol (DTT) AT 56°C for 45 min to break disulphide bonds and alkylated with 55 mM idoacetamide to prevent disulphide reformation. Tryptic digestion was performed overnight using sequencing grade trypsin (15ng/µL) in 25mM ABC containing 1 mM cacl_2._ Peptides were sequentially extracted with 5% formic acid in ACN and desalted using C18 spin columns. The desalted peptides were analysed on a Q-Exactive Orbitrap mass spectrometer (Thermo Fisher Scientific). Raw MS data from the three sample batches were processed using Proteome Discoverer v2.2.0388 with Sequest HT search against a curated human cornea specific UniProtKB FASTA database ensuring tissue specific protein identification. Label Free Quantification (LFQ) was employed to compare protein abundance across the three batches without isotopic labelling, enabling the assessment of differential extracellular matrix (ECM) protein expression. Protein identification were stringently filtered using the following criteria: FDR ≤ 1% (99% confidence), ≥1 unique peptides, ≥2 total peptides, sequest HT score ≥1.00, and master proteins were retained. Commonly identifies proteins across the three sample sets were visualised using interactiVenn.

### Network Pathway Analysis

Network pathway analysis was performed on the list of proteins obtained through proteomic analysis. The differentially expressed proteins were analysed for the association in canonical pathways, disease & disorders, molecular & cellular functions and physiological system development & functions using ingenuity pathway analysis (IPA) software.

## Results & Discussion

### Corneal Extra cellular Matrix Proteome analysis

Based on the proteomic analysis, an average of 18 proteins were identified in all the three sample sets as shown in **Table 1, Table 2 & Table 3** respectively. Among the identified proteins, extra cellular matrix proteins, glycoproteins, and peptidoglycans were detected by cross-referencing the results with list of proteins (Appendix I) available in the human cornea database https://www.uniprot.org/. The identified proteins (Appendix III, IV & V) were subsequently analysed and filtered using specific parameters using Thermo Proteome Discoverer software (version 2.2.0.388). The proteins were filtered based on Protein FDR confidence which is false discovery date, a statistical measure used to estimate the proportion of incorrect (false positive) identifications among all positive identifications. This is especially important when matching MS/MS spectra to peptides or proteins. A low FDR (FDR ≤ 0.01) is desirable as used in the current study, indicating a minimal number of false positives, thereby enhancing the reliability and confidence in the identified peptides and proteins. An FDR of 0.01 implies a 99% confidence that the identified peptide/protein is not a false positive. Following this, filtration was performed based on unique peptides, which is the total number of peptide sequences unique to the protein groups. The threshold is set to greater than or equal to 1 indicating the protein identified must have at least one peptide and not shared with any other protein which ensures specificity. Additionally, proteins were filtered based on the total number of distinct peptides identified from MS/MS spectra. The filter is set to greater than or equal to 2 peptides (≥2 peptides) meaning that only proteins identified by at least two different peptide sequences will be included in the results. Sequest HT (High Throughput) is used for peptide identification in MS-based proteomics. It compares experimental MS/MS spectra to theoretical spectra of peptides generated *in silico* from a protein database. This score reflects the quality of the match between the observed spectrum and the predicted fragment ion spectrum of a peptide. The higher score, the better match, and the higher confidence in peptide identification. Based on this, the filter is set to greater than or equal to 1.00 and the proteins were filtered. Additionally, only the master proteins, the primary representative proteins within each protein group were retained for analysis. Using all these filtering criteria, the common proteins shared among all three sample sets were identified using InteractiVenn (Bioinformatics 16:169), a tool for visualizing overlaps between datasets.

**Table 1.**
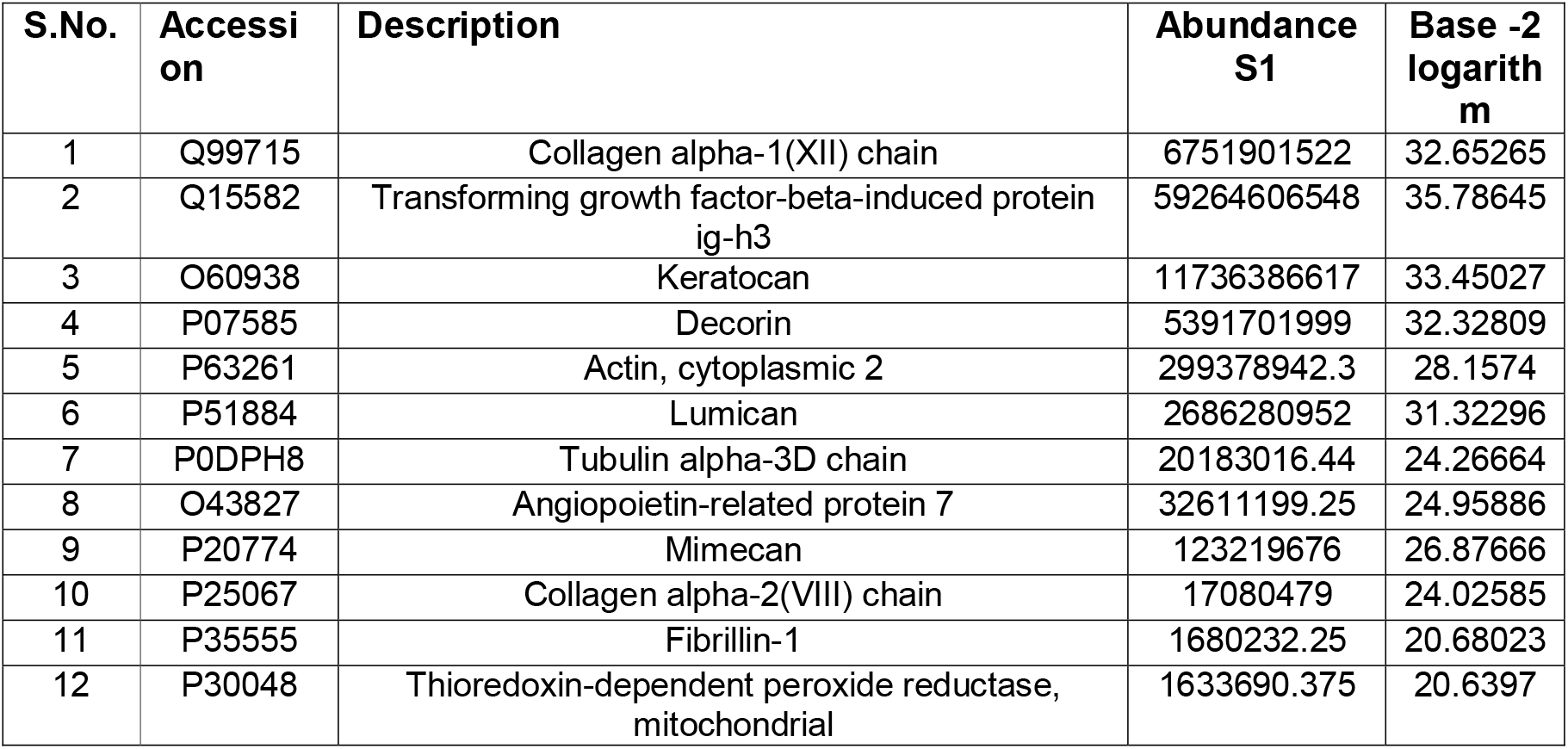
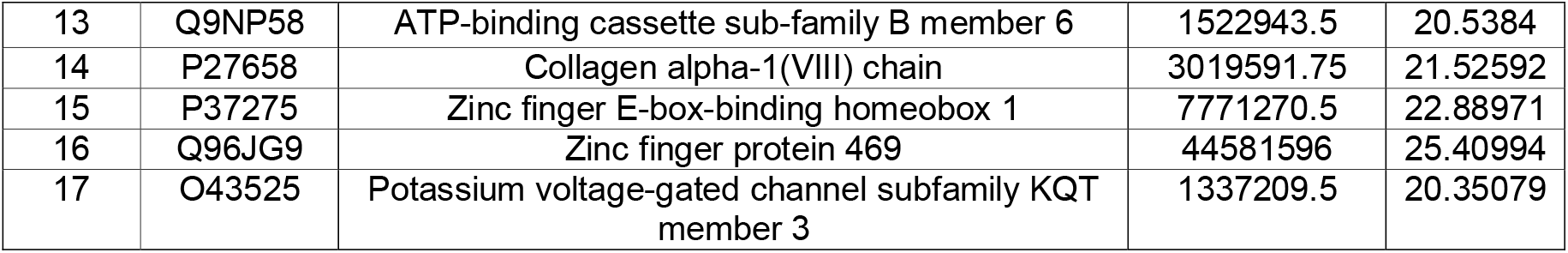
List of proteins identified in sample set 1 and compared based on the abundance and respective base-2 logarithm.

**Table 2.** List of proteins identified in sample set 2 and compared based on the abundance and respective base-2 logarithm.

| S.No. | Accession | Description | Abundance S2 | Base -2 logarithm |
| --- | --- | --- | --- | --- |
| 1 | Q99715 | Collagen alpha-1(XII) chain | 8171699423 | 32.92799 |
| 2 | Q15582 | Transforming factor-beta-induced protein ig-h3 | 50691739282 | 35.56103 |
| 3 | O60938 | Keratocan | 13366356436 | 33.63789 |
| 4 | P07585 | Decorin | 6252056423 | 32.54168 |
| 5 | P51884 | Lumican | 2751509633 | 31.35758 |
| 6 | P63261 | Actin, cytoplasmic 2 | 284375956.8 | 28.08322 |
| 7 | P0DPH8 | Tubulin alpha-3D chain | 19441356.69 | 24.21263 |
| 8 | P20774 | Mimecan | 128318620 | 26.93516 |
| 9 | O43827 | Angiopoietin-related protein 7 | 46373053 | 25.46678 |
| 10 | P35555 | Fibrillin-1 | 12405322.38 | 23.56446 |
| 11 | P25067 | Collagen alpha-2(VIII) chain | 14527961.5 | 23.79233 |
| 12 | P30048 | Thioredoxin-dependent peroxide reductase, mitochondrial | 1503748.25 | 20.52013 |
| 13 | P27658 | Collagen alpha-1(VIII) chain | 17935270 | 24.0963 |
| 14 | Q7Z7G0 | Target of Nesh-SH3 | 586956 | 19.16289 |
| 15 | Q17RW2 | Collagen alpha-1(XXIV) chain | - | - |
| 16 | Q9UQC9 | Calcium-activated chloride channel regulator 2 | 32888750 | 24.97109 |
| 17 | Q9HCD6 | Protein TANC2 | 24609242 | 24.55269 |
| 18 | Q9UMD9 | Collagen alpha-1(XVII) chain | 11353593 | 23.43664 |

**Table 3.** List of proteins identified in sample set 3 and compared based on the abundance and respective base-2 logarithm.

| S.No. | Accession | Description | Abundance S3 | Base -2 logarithm |
| --- | --- | --- | --- | --- |
| 1 | Q99715 | Collagen alpha-1(XII) chain | 8359338706 | 32.96074 |
| 2 | Q15582 | Transforming growth factor-beta-induced protein ig-h3 | 55521851858 | 35.69234 |
| 3 | P07585 | Decorin | 7355561950 | 32.77619 |
| 4 | O60938 | Keratocan | 17366245189 | 34.01557 |
| 5 | P63261 | Actin, cytoplasmic 2 | 326765844.3 | 28.28368 |
| 6 | P51884 | Lumican | 3572648380 | 31.73435 |
| 7 | O43827 | Angiopoietin-related protein 7 | 52962208.5 | 25.65846 |
| 8 | P20774 | Mimecan | 222491805 | 27.72918 |
| 9 | P0DPH8 | Tubulin alpha-3D chain | 18201650.88 | 24.11757 |
| 10 | P27658 | Collagen alpha-1(VIII) chain | 25367061.88 | 24.59645 |
| 11 | P35555 | Fibrillin-1 | 6433240.75 | 22.61711 |
| 12 | P25067 | Collagen alpha-2(VIII) chain | 19736570.5 | 24.23437 |
| 13 | P30048 | Thioredoxin-dependent peroxide reductase, mitochondrial | 2059120.25 | 20.9736 |
| 14 | Q9UMD9 | Collagen alpha-1(XVII) chain | 41404185.66 | 25.30327 |
| 15 | Q7Z7G0 | Target of Nesh-SH3 | 5037841.875 | 22.26437 |
| 16 | P13796 | Plastin-2 | 1150999.875 | 20.13445 |
| 17 | Q96JG9 | Zinc finger protein 469 | 1395024.125 | 20.41185 |
| 18 | P35968 | Vascular endothelial growth factor receptor 2 | 45249752 | 25.43140 |
| 19 | Q9Y2V3 | Retinal homeobox protein Rx | 6124316 | 22.54611 |

Consequently, proteins beyond the 13 commonly identified were excluded as they did not meet the criteria set by the applied filter. The unidentified/low abundance proteins were ATP-binding cassette sub-family B member 6, Zinc finger E-box-binding homeobox 1, Zinc finger protein 469, Potassium voltage-gated channel subfamily KQT member 3, Vascular endothelial growth factor receptor 2, Calcium-activated chloride channel regulator 2, Protein TANC2, Retinal homeobox protein Rx, Plastin-2, Collagen alpha-1(VIII) chain, Collagen alpha-1(XVII) chain, Collagen alpha-1(XXIV) chain, and Target of Nesh-SH3. The excluded proteins represent a diverse group with critical roles in cell signalling, ECM organization, tissue development, and structural maintenance, particularly in the cornea, nervous system, and epithelial tissues. Their loss during proteomic filtering may reflect low abundance, complex structure, or limited peptide detectability, rather than a lack of biological importance. Their shared involvement in developmental pathways, cellular architecture, and disease associations underscores the need for tailored approaches in proteomic analysis to retain such functionally significant proteins.

A total of 13 proteins which are majorly extra cellular matrix proteins were common in all the three sample sets which have a prominent role in proliferation and remodelling of cornea, concerning wound healing and regeneration. **Fig 1** represents the number of identified proteins in the individual sample and number of proteins that are common among the two sample sets and the common proteins among all the three groups. Based on the common proteins identified, the proteins were analysed and categorized graphically comparing the base-2 logarithm of abundance of all the proteins in each sample set as shown in **Fig 2**, demonstrates the relative levels of each protein, and pinpointing those with comparable amounts in all samples (**Table 4)**.

**Table 4.** List of common proteins identified from three sample sets and compared based on the abundance (log2).

| S.No | Accession | Description | Symbol | Abundance S1 | Abundance S2 | Abundance S3 |
| --- | --- | --- | --- | --- | --- | --- |
| 1 | P30048 | Thioredoxin-dependent peroxide reductase, mitochondrial | PRDX3 | 20.6397 | 20.52013 | 20.9736 |
| 2 | P35555 | Fibrillin-1 | FBN1 | 20.68023 | 23.56446 | 22.61711 |
| 3 | P27658 | Collagen alpha-1(VIII) chain | COL8A1 | 21.52592 | 24.0963 | 24.59645 |
| 4 | P25067 | Collagen alpha-2(VIII) chain | COL8A2 | 24.02585 | 24.79233 | 24.23437 |
| 5 | P0DPH8 | Tubulin alpha-3D chain | TUBA3D | 24.26664 | 24.21263 | 24.11757 |
| 6 | O43827 | Angiopoietin-related protein 7 | ANGPTL7 | 24.95886 | 25.46678 | 25.65846 |
| 7 | P20774 | Mimecan | OGN | 26.87666 | 26.93516 | 27.72918 |
| 8 | P63261 | Actin, cytoplasmic 2 | ACTG1 | 28.1574 | 28.08322 | 28.28368 |
| 9 | P51884 | Lumican | LUM | 31.32296 | 31.35758 | 31.73435 |
| 10 | P07585 | Decorin | DCN | 32.32809 | 32.54168 | 32.77619 |
| 11 | Q99715 | Collagen alpha-1(XII) chain | COL12A1 | 32.65265 | 32.92799 | 32.96074 |
| 12 | O60938 | Keratocan | KERA | 33.45027 | 33.63789 | 34.01557 |
| 13 | Q15582 | Transforming growth factor-beta-induced protein ig-h3 | TGFBI | 35.78645 | 35.56103 | 35.69234 |

**Fig 1.**
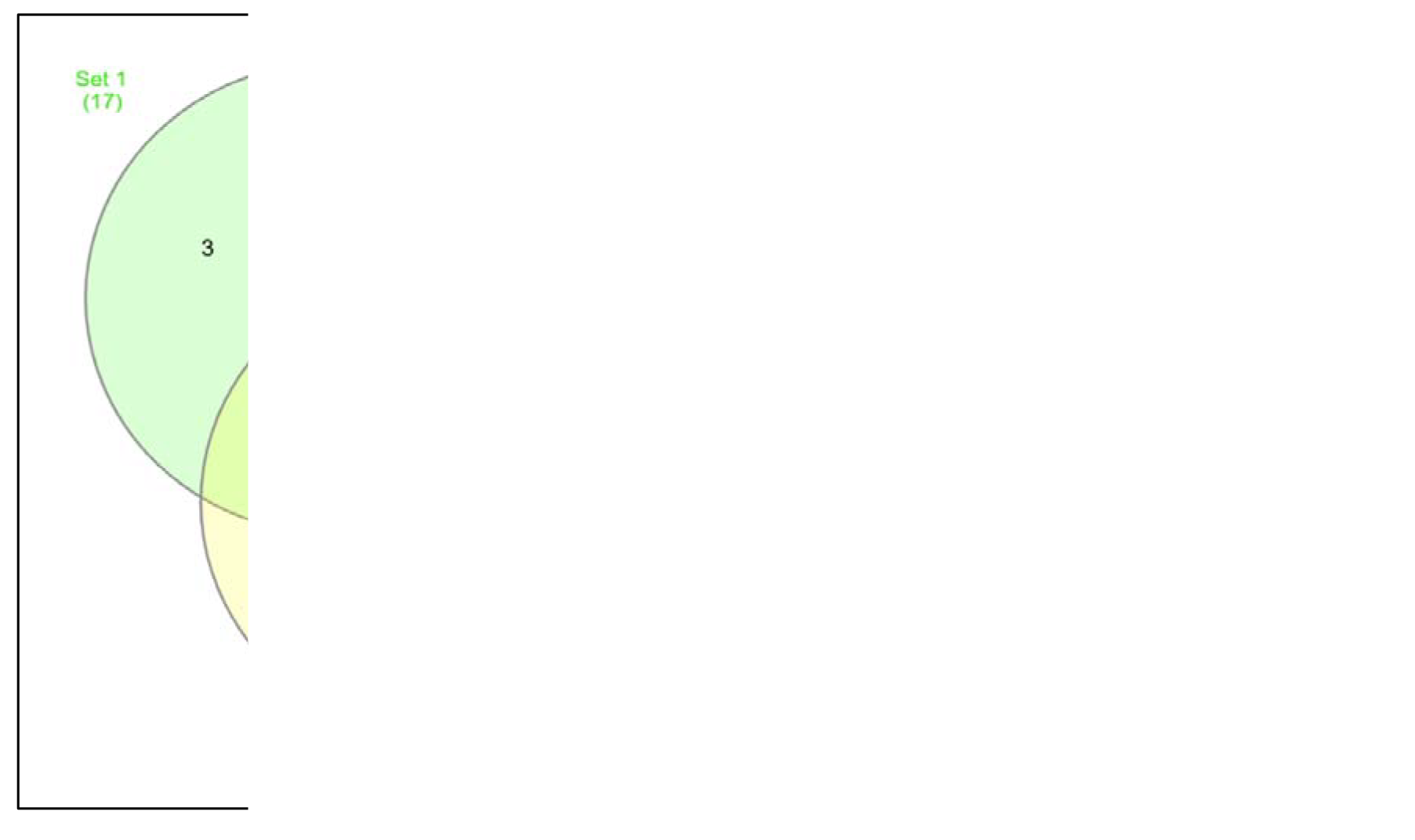
Venn diagram of identified proteins; S1-17 proteins ; S2-18 proteins; S3-19 proteins; Common proteins identified among the three samples-13 proteins

**Fig 2.**
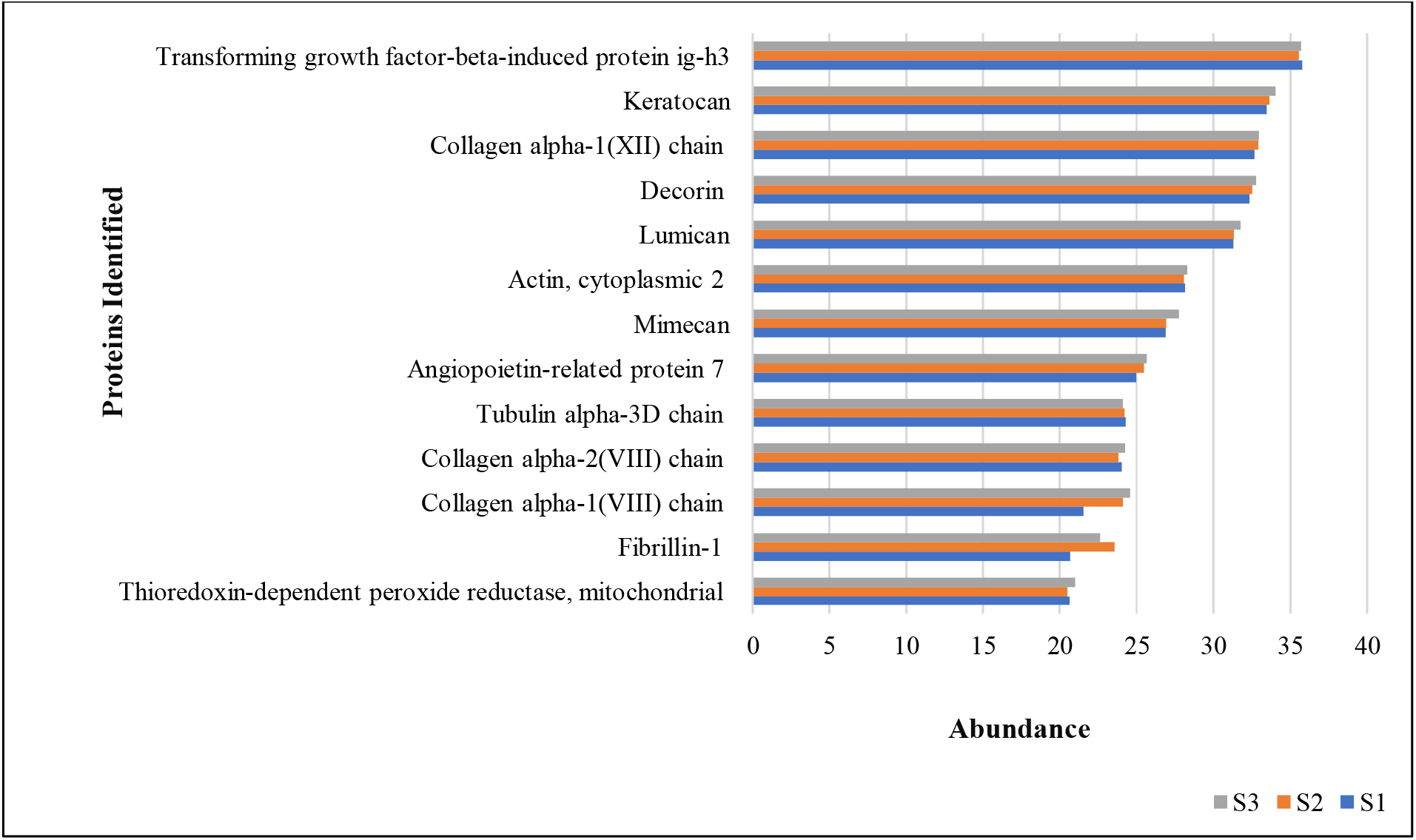
Common proteins identified were compared based on the abundance obtained from three sample sets (n=3)

The 13 key extracellular matrix (ECM) proteins identified across the three samples play critical roles in corneal regeneration and wound healing by regulating collagen organization, stromal remodelling, inflammation modulation, and growth factor signalling, functions central to maintaining corneal transparency and structural integrity.

The predominant form of collagen XII is present in human stroma and sclera that regulates lamellar organization and is expressed in regions of high mechanical stress. During stromal injury, collagen XII is upregulated and is present in scarred stroma indicating its role in stromal regeneration, wound healing, and remodelling. The decrease in collagen XII as well as proteoglycans and collagen I deposition observed in keratoconus corneas^3^.

The TGF-β in the corneal stroma is significantly regulated by collagen XII and this activity controls matrix deposition in the stromal structure during development and homeostasis. The mRNA expression studies of *Col12a1* showed stable level in the mature cornea and it increased to maximum during maturation^9^.

TGF- β is a multifunctional signalling molecule that regulates diverse cellular responses^10^. In the healthy cornea, TGF- β1 is generally present in corneal epithelial cells while TGF- β2 and TGF- β3 were identified in the extracellular environment^11^. In epithelial wounds, TGF- β stimulates the migration and proliferation of corneal epithelial cells and activates keratocyte proliferation and myofibroblast differentiation. However, high levels of TGF- β causes aberrant collagen deposition resulting in corneal scarring and visual impairment. Therefore, activation and regulation of TGF- β are crucial for sustaining corneal clarity and vision post injury^10^.

In the cornea, TGF- β contributes to the regulation of inflammation in the cornea and promote immunotolerance post-injury by promoting the recruitment and activation of T-cells. Furthermore, it has been identified that TGF-β stimulates the synthesis of lipoxins, which are anti-inflammatory proteins that aid in tissue repair and inflammation resolution^10^.

Decorin, a small leucine-rich proteoglycan (SLRP), found in collagen fibrils, that make up the extracellular matrix of all connective tissues. It plays a major role in regulating collagen fibrillogenesis, ECM production, cell cycle progression by interacting with epidermal growth factor (EGF), tumor necrosis factor alpha (TNF α), transforming growth factor beta (TGF- β), fibronectin, and various other collagens^12^.

Decorin regulates the spacing and alignment of collagen fibrils in the cornea, which is crucial for its transparency and for corneal healing after injury.

In TGF-β regulation, decorin binds to transforming growth factor-beta, (TGF-β), a cytokine involved in fibrotic tissue formation. Decorin promotes healthy corneal regeneration by limiting the excessive deposition of extracellular matrix (ECM) and fibrosis by concealing TGF-β. By regulating TGF-β, decorin helps to prevent this differentiation, reducing scar tissue formation in the cornea.

During corneal wound healing, decorin interacts with inflammatory cytokines like interleukin-1 (IL-1) and tumour necrosis factor-alpha (TNF-α), reducing their effects and preventing chronic inflammation^13^.

Lumican is a critical regulator of the extracellular matrix, particularly influencing collagen organization. In the cornea, it functions primarily as a keratan sulphate proteoglycan. This role is essential for maintaining corneal transparency by regulating the assembly of collagen fibres in the stromal layer. In the corneal stroma, lumican is expressed by stromal keratocytes under normal, unwounded conditions which are responsible for maintaining the integrity and clarity of the cornea by secreting components like collagen and lumican.

In the context of corneal injury, lumican is involved in regulating both the inflammatory response and the subsequent tissue remodelling that is essential for proper healing. It can influence the infiltration of immune cells, such as neutrophils and macrophages, into the wounded area^14^.

Keratan sulphate (KS), is a type of glycosaminoglycan that was initially discovered in the cornea^15^.

Keratan sulphate proteoglycan plays a key role in the transparency of cornea. It is present in large quantities in the corneal stroma, that regulates the organization of collagen fibrils and is involved in wound repair of epithelial tissues of the human cornea^16^.

Osteoglycin also known as mimecan is a class III SLRP. It is primarily located in the corneal stroma, with some expression in the corneal epithelium and epithelial basement membrane. The function of osteoglycin in the cornea is likely intertwined with the actions of other SLRPs like lumican, keratocan, and decorin, which also play crucial roles in maintaining corneal structure. These proteins which are known to interact with collagen fibrils and help maintain their proper spacing and organization within the corneal stroma, contributing to corneal transparency^17,18^.

Actin Cytoplasmic 2, serving as a critical component of the cell’s cytoskeleton, is essential for corneal function. This protein supports the structure of corneal epithelial cells, enables their migration during wound healing and maintains the epithelial barrier^19^.

### Network Pathway Analysis

From the list of proteins identified, it was found that Integrin cell surface interactions, collagen chain trimerization, assembly of collagen fibrils, and other multimeric structures, collagen degradation and collagen biosynthesis and modifying enzymes are the most highly associated canonical pathways (Table 5a). Hereditary Disorder, ophthalmic disease, organismal injury, and abnormalities, developmental disorder and cancer are the most associated diseases and disorders (Table 5b). The major molecular and cellular functions associated with the regeneration process are, cellular movement and cell morphology, cellular function and maintenance, cell-to-cell signalling and interaction and lipid metabolism (Table 5c). The physiological system development and function associated with regeneration based on analysis are visual and cardiovascular system development and function, hair and skin, connective tissue development and function and morphology (Table 5d).

**Table 5.** List of top five canonical pathways, disease and disorders and molecular and cellular functions involved based on network pathway analysis.

| Name | p-value |
| --- | --- |
| **a. Top Canonical Pathways** |  |
| Integrin cell surface interactions | 3.26E-08 |
| Collagen chain trimerization | 1.50E-07 |
| Assembly of collagen fibrils and other multimeric structures | 5.98E-07 |
| Collagen degradation | 6.90E-07 |
| Collagen biosynthesis and modifying enzymes | 8.31E-07 |
| **b. Diseases and Disorders:** |  |
| Hereditary Disorder | 3.19E-02- 5.85E-21 |
| Ophthalmic Disease | 3.03E-02- 5.85E-21 |
| Organismal Injury and Abnormalities | 3.45E-02- 5.85E-21 |
| Developmental Disorder | 3.23E-02- 9.07E-17 |
| Cancer | 3.45E-02- 9.10E-08 |
| **c. Molecular and Cell Functions:** |  |
| Cellular Movement | 2.23E-02- 1.02E-04 |
| Cell Morphology | 2.89E-02- 1.87E-04 |
| Cellular Function and Maintenance | 2.89E-02- 1.87E-04 |
| Cell-To-Cell Signalling and Interaction | 3.05E-02- 2.04E-04 |
| Lipid Metabolism | 1.85E-02- 4.94E-04 |
| **d. Physiological System Development and Function** |  |
| Visual System Development and Function | 5.47E-03 - 5.84E-05 |
| Cardiovascular System Development and Function | 3.45E-02 - 2.04E-04 |
| Hair and Skin Development and Function | 2.15E-02 - 1.03E-03 |
| Connective Tissue Development and Function | 1.09E-02 - 1.10E-03 |
| Organ Morphology | 6.56E-03 - 1.10E-03 |

The p-values obtained from this pathway analysis is a statistical measure used to determine a very strong significance of the association between the input dataset (the identified proteins in the given samples) and a particular canonical pathway or network. In this study, the p-value of 8.31E-07 for the pathway “collagen biosynthesis and modifying enzymes” indicates a highly significant correlation between the identified ECM proteins and the canonical pathway, followed by collagen degradation and assembly of collagen fibrils and other multimeric structures, integrin cell surface interactions and others as listed in table 5.

Network pathway analysis highlighted several key biological processes, diseases, and functional associations relevant to the cornea. Among the top canonical pathways Integrin cell surface interactions” showed the most significant association (*p* = 3.26 × 10^−^□, followed by Collagen chain trimerization (*p* = 1.50 10^−^□, Assembly of collagen fibrils and other multimeric structures (*p* = 5.98 × 10^−^□, “Collagen degradation” (*p* = 6.90 × 10^−^□, and Collagen biosynthesis and modifying enzymes (*p* = 8.31 × 10^−^□, emphasizing the central role of collagen metabolism and integrin-mediated signaling in corneal structure and function ^20,21^. Disease and disorder associations included hereditary disorders, ophthalmic diseases, organismal injury and abnormalities, developmental disorders, and cancer, all with highly significant p-values, indicating that alterations in these pathways could contribute to a wide range of pathological outcomes. Molecular and cellular functions were linked to cellular movement, morphology, maintenance, cell-to-cell signalling, and lipid metabolism, suggesting their involvement in corneal repair and homeostasis ^22,23^. Furthermore, physiological system development and function analysis revealed strong associations with visual system development, as well as cardiovascular, hair and skin, and connective tissue development, alongside organ morphology, reflecting the interconnected nature of ECM-related pathways in maintaining tissue integrity and specialized function.

The genes and proteins identified through data analysis using IPA software as mentioned in Appendix II encode few ECM proteins. Mutations or alterations in these genes can disrupt corneal structure, integrity, function, and overall maintenance, potentially leading to disease.

## Conclusion

Proteomic profiling of decellularised human corneal ECM (hdECM) identified 13 key extracellular matrix proteins conserved across all donar samples. These proteins including collagen XII, decorin, lumican and keratan sulphate sulphate proteoglycans are central to stromal organization, transperancy, and wound healing. High confidence filtering revealed a robust ECM framework that supports corneal regeneration and highlights hdECM’s potential in therapeutic and tissue engineering applications.

## Summary

What was known before:

- The complete proteomic profile of the human cornea is available at on https://www.uniprot.org/.
- However, the specific data on extra cellular matrix proteins isolated from decellularized human corneas has not been reported yet.

What this study adds:

- To best of our knowledge, this is the first study to identify extracellular matrix proteins from lyophilized sample prepared from decellularized bio banked human corneas.

## Data availability

The proteomic data obtained from the study has been deposited to the ProteomeXchange consortium via the PRIDE21 partner repository with the dataset Accession No. PXD060268

## Author Contributions

**Khyathi Ratna Padala** performed proteomic analysis and data interpretation. **PV Anusha** helped in proteomics study., **Mohammed M Idris** helped in proteomics study and network pathway analysis, **Vivek Singh** participated in proof reading of the manuscript. **Shibu Chameettachal** participated in proof reading of the manuscript. **Kiran Kumar Bokara** conceptualised, supervision, manuscript editing and proof reading of the manuscript

